# Propensity-Adjusted AUC under Nuisance-Model Misspecification

**DOI:** 10.64898/2026.11.11.765682

**Authors:** Caleb Heinzman, Yi Shang, Huazhang Guo, Ye Duan, Mai He

## Abstract

Propensity adjustment compares medical-image detector scores within groups with similar measured properties, such as stain or focus. We examine its sensitivity to the nuisance model that forms those groups. Under Gaussian latent-index assumptions and exact linear-index strata, a closed-form correlation gives an exact boundary for adjusted AUC to fall below one half. Quadrature matches 36 simulation cells within 0.008. Inflation and attenuation relative to correct adjustment are empirical findings, not general ordering results. An overlap screen and a self-stratified control introduced for this study miss 140 of 141 simulated cells with absolute error above 0.05: checking the fitted propensity cannot certify completeness of the adjustment set. On real pathology corpora, logistic and boosted-tree models using the same 16 features disagree. Among splits where both pass both checks, mean adjusted-AUC gaps are 0.057 on acid-fast bacilli (AFB), 0.048 on Kaggle, and 0.049 on PCam, over three, five, and three splits respectively. These gaps measure model sensitivity; neither estimate is known to be correct. A controlled colour shift moves adjusted AUC relative to model-specific references in the predicted direction in 78 of 80 label-dependent cells. Broader balance and label-fit diagnostics favour trees without certifying their adjusted AUC. We recommend reporting sensitivity across nuisance models alongside these diagnostics. Reversal relative to known truth occurs in constructed stress tests; its frequency in practice remains unknown.

## 1 Introduction

A medical-image detector can get a high area under the receiver operating characteristic curve (ROC-AUC, or AUC below) for the wrong reason (Geirhos et al., 2020; DeGrave et al., 2021; Zech et al., 2018; Badgeley et al., 2019; Oakden-Rayner et al., 2020). Stain, focus and field crowding can predict labels through collection artifacts that do not transfer to another scanner or patient mix. We call these unwanted sources of label information nuisance factors. Which properties are nuisance is a clinical judgment, discussed in §2 and §6.

Covariate-adjusted ROC analysis compares positives and negatives with similar measured properties, then averages their discrimination (Janes & Pepe, 2009; Pepe et al., 2013; Janes et al., 2009). Related propensity approaches (Galadima & McClish, 2019; Mushfiquee & Rahman, 2022) motivate a scalar summary. Here a nuisance model predicts the label from measured stain, focus and texture features. We bin images by its predicted probability and average within-bin detector AUCs, weighted by the number of positive–negative pairs. Correct adjustment requires the nuisance model to represent the relevant properties and their functional form. We study what happens when it does not, and whether checks of its own inputs and outputs can reveal the error. Our intended readers are evaluators who report shortcut-adjusted AUCs for imaging detectors.

The ingredients are established. Omitted-variable bias, balancing scores and Simpson-type reversal under partial adjustment are standard in causal inference, which has long warned, for treatment effects, that balance on the properties used says nothing about those left out (Weitzen et al., 2005; Austin, 2011; Ho et al., 2007; Imai et al., 2008; Westreich et al., 2011; Sant’Anna & Song, 2019); negative-control designs (Lipsitch et al., 2010) and over-adjustment analyses (Schisterman et al., 2009) address related failures. Galadima & McClish (2019) studied bias from propensity adjustment when AUC measures the effect of a binary exposure on a continuous outcome, a different application from diagnostic discrimination. Our contribution is to work out what these mechanisms imply for the adjusted AUC and its checks. We organise the paper around three findings.

### *An exact reversal boundary* (§3)

In a Gaussian latent-index model with exact linear-index strata, a closed-form correlation determines the sign of adjusted AUC minus one half (Theorem 1). Quadrature and simulations also describe inflation and attenuation relative to correct adjustment in the designs studied. Those comparisons are empirical; the theorem concerns reversal relative to chance.

### *The checks’ blind spot* (§4)

For a model that is right on the properties it uses, conditioning on its exact propensity balances those properties whatever it left out, and a score stratified on itself has AUC one half by definition (Proposition 6, Lemma 5). In finite-bin simulations, large errors almost always pass the overlap screen and the self-stratified control. These two checks cannot certify that nothing was omitted; a wrong functional form on the properties used is a different failure, which more flexible diagnostics can detect (§5).

### *Sensitivity to nuisance-model choice* (§5)

On real pathology corpora, logistic and boosted-tree models using the same 16 features differ by about 0.05. The discrepancy persists on AFB, Kaggle and PCam when restricted to splits where both models pass both checks. Different fitted propensities change the conditioning groups and pair weights, so these gaps do not identify either model’s error. Broader fit diagnostics favour trees, while the available data provide no known correct adjusted AUC. We therefore report both estimates and give a checklist for assessing sensitivity.

## 2 Setup: adjusted AUC, the nuisance model, and the checks

An image *X* has a binary label *Y* and a detector score *S*(*X*). A vector *Z* of measured image properties (color, focus, texture, edge, frequency, and morphology statistics; Appendix C) is the nuisance descriptor. The nuisance model estimates *e*(*Z*) = *P* (*Y* =1 | *Z*), and its fitted value *ê* is the propensity.

To compute the adjusted AUC we sort the test images by *ê* into *q* bins of equal count (quantile strata; *q* = 20 at our test-fold sizes), compute the detector’s AUC within each bin, and average the bin AUCs with weights proportional to the number of positive–negative pairs in the bin. In population form, write *B* = *ê*(*Z*), *F*_*B*_ for its distribution, *π*(*b*) = *P* (*Y* =1 | *B*=*b*), and *S*_1_, *S*_0_ for the scores of a positive and a negative image. The within-stratum AUC at propensity *b* is 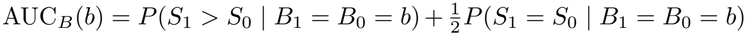 and the adjusted AUC is

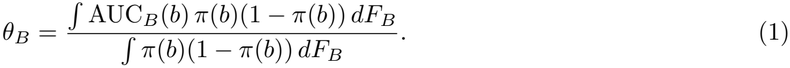

For a continuous propensity this is the population target of increasingly fine, equal-mass bins; a bin with no positive–negative pairs contributes nothing. With a fixed number *q* of bins, the implemented estimator targets *θ*_*q*_ = ∑_*k*_ *w*_*k*_ AUC_*k*_*/*∑_*k*_ *w*_*k*_, where bin *k* has probability mass *p*_*k*_, positive share *π*_*k*_, within-bin AUC AUC_*k*_, and pair weight 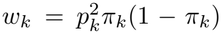. With equal masses this is the *q*-bin analogue of (1); tied propensities, which must share a bin, can make masses unequal and so change the weights as well as the conditioning. In our real-data comparisons the fitted propensities have essentially no ties (bin sizes differ by at most 1.3%), and weighting bins by *π*_*k*_(1 *π*_*k*_) instead, or breaking ties at random, changes any split’s adjusted AUC by at most 0.0002 (Appendix G). The true propensity *e*(*Z*) is a balancing score (Rosenbaum & Rubin, 1983); an estimated propensity, or finite bins of it, need not balance *Z*.

### Related estimands

Janes & Pepe (2009) average covariate-specific ROC curves over the covariate distribution of the cases; subgroup AUC is computed directly within predefined groups, with no fitted propensity. Our target stratifies on a fitted propensity and weights strata by positive–negative pairs (overlap weights). The three estimate different quantities, and our results are established only for (1). Even with the correct nuisance model, *θ*_*B*_ conditions only on the propensity, a one-dimensional summary of *Z*: it compares positives and negatives that are equally likely to be positive given their properties, not positives and negatives with equal properties. It is therefore neither discrimination conditional on the full vector *Z* nor an intrinsic measure of how well the detector recognises disease; it measures the detector’s discrimination beyond what the measured properties predict about the label.

### Checks

We introduce two checks for this study to examine what can be learned from the fitted propensity alone. The overlap screen flags the result if more than 20% of strata contain one label only. The self-stratified control takes the fitted propensity itself as the score, computes its adjusted AUC within the same strata with no refitting, and passes when the value lies in [0.45, 0.55]. This is a self-consistency diagnostic for the binning procedure: in exact propensity strata the score is constant, so its AUC is one half. With finite bins, deviations can reveal residual label gradients or limited overlap. A pass cannot establish that relevant covariates were included. This pair of checks and its cutoffs are study conventions, not an established certification procedure. Their limited scope motivates the comparison with broader balance and specification diagnostics in §5; Appendix C examines cutoff sensitivity.

### Nuisance models and detectors

C1 is logistic regression on 16 color and focus features. C2 is gradient-boosted trees on the same 16. C3 is boosted trees on 62 color, focus, texture, and edge features that include the 16. C4 is boosted trees on 95 features that add 18 frequency and 15 morphology features. We call logistic models the linear class and tree ensembles the tree class. The main detector is the zero-shot prompt-ensemble score of a vision–language model (Zhang et al., 2023). A self-trained student of it (a logistic model on the same frozen embeddings, fitted to the prompt score’s own top and bottom 20% as pseudo-labels) and supervised probes trained on the real labels appear in Appendix F.

### Terms

*Families I–III* are the Gaussian simulation designs of §3–4. *Groups A and B* are descriptor subsets held in or out of the nuisance model by the held-out tests (§5). *Truth* is the adjusted AUC under the correct nuisance model, known only when we generate the labels; an absolute error above 0.05 is called *material*.

### What counts as nuisance

The theory assumes the hidden disease signal is independent of the measured properties. In pathology, texture and morphology may partly reflect the organism itself, and adjusting for them then removes real signal as well as shortcuts. A lower adjusted AUC is therefore not necessarily a more valid one; the choice of adjustment set needs clinical input, and results should be reported with and without properties that may carry disease signal.

## 3 Reversal theorem and empirical adjustment effects

A nuisance model can omit a measured feature or fail to represent a combination of features. That direction then varies within its strata, and detector discrimination can reflect both the target signal and the uncontrolled direction.

We distinguish two comparisons. *Reversal* means that adjusted AUC falls below chance, one half; Theorem 1 gives its exact boundary in a Gaussian model. *Inflation* and *attenuation* compare adjusted AUC with the value under correct nuisance adjustment. In the designs studied, alignment of the detector with the missed label direction raises the estimate, while ignoring that direction lowers it. These latter patterns are supported by quadrature and simulation, not by a general ordering theorem. In synthetic experiments the reference is known from the generating model; on real labels it is unknown.

### The two-property example

Take two measured properties *Z*_1_ and *Z*_2_, a hidden target *U*, and a detector noise term, all standard Gaussian. The label’s log-odds are *α*(*Z*_1_ + *Z*_2_) + *U*, so the label loads *α* on each property. The detector score is *U* + *λ*(*Z*_1_ − *Z*_2_) plus noise: it carries the target with loading one and loads −*λ* on *Z*_2_, against the label. The nuisance model is logistic regression on *Z*_1_ alone, so *Z*_2_ is the missed direction, and inside its strata positives have larger *Z*_2_ than negatives.

Varying *λ* illustrates these effects (Table 1, Family I, *α* = 1, law columns). At *λ* = 0 the detector ignores *Z*_2_ and the estimate is 0.648 against a truth of 0.670, an attenuation of 0.022. At *λ* = 1 the loading against *Z*_2_ cancels the target signal exactly and the estimate is 0.500, equal to chance. At *λ* = 2 the estimate is 0.415 against a truth of 0.575, reversing the ranking within strata. The estimate is below one half exactly when *αλ >* 1.

**Table 1:** The formula against simulation (adjusted AUCs). Rows are settings of the label loading *α* and the detector loading *λ*. “Truth” is the adjusted AUC under the correct nuisance model; “wrong model” under the misspecified one. “Law” is predicted by (2) and quadrature with no simulation; “sim” is the mean over ten simulated replicates. Family I is the two-property example (the detector’s loading on the missed property opposes the label’s, so *λ >* 0 lowers the estimate). Family III has twelve correlated properties (AR(1), correlation 0.6) and a wrong model fit on six; its law columns use the ideal subset index and its simulation columns use fitted boosted trees; the hidden-column loading has the label’s sign, so *λ >* 0 inflates (Appendix B). Bold entries indicate adjusted AUCs below 0.50.

|  |  | Family I (omitted *Z*<sub>2</sub>) |  |  |  | Family III (6 of 12 used) |  |  |  |
|---|---|---|---|---|---|---|---|---|---|
|  |  | truth |  | wrong model |  | truth |  | wrong model |  |
| *α* | *λ* | law | sim | law | sim | law | sim | law | sim |
| 0.5 | 0 | 0.668 | 0.668 | 0.662 | 0.662 | 0.668 | 0.669 | 0.665 | 0.667 |
| 0.5 | 1 | 0.618 | 0.620 | 0.565 | 0.563 | 0.653 | 0.653 | 0.675 | 0.681 |
| 0.5 | 2 | 0.574 | 0.574 | 0.500 | **0.498** | 0.624 | 0.625 | 0.653 | 0.660 |
| 1 | 0 | 0.670 | 0.672 | 0.648 | 0.650 | 0.669 | 0.668 | 0.660 | 0.655 |
| 1 | 1 | 0.619 | 0.619 | 0.500 | 0.500 | 0.654 | 0.653 | 0.708 | 0.711 |
| 1 | 2 | 0.575 | 0.575 | **0.415** | **0.416** | 0.625 | 0.626 | 0.703 | 0.708 |
| 2 | 0 | 0.671 | 0.671 | 0.617 | 0.620 | 0.671 | 0.672 | 0.643 | 0.642 |
| 2 | 1 | 0.621 | 0.622 | **0.404** | **0.409** | 0.656 | 0.655 | 0.755 | 0.755 |
| 2 | 2 | 0.576 | 0.577 | **0.296** | **0.293** | 0.627 | 0.626 | 0.780 | 0.782 |

### Exact statement

The example is a case of a Gaussian latent-index model: *Z* ∼ *N* (0, Σ), a target latent *U* ∼ *N* (0, 1), noise *ε* ∼ *N* (0, *σ*^2^), all independent, with Σ positive definite, *σ*^2^ *>* 0, and *a, w* = 0; label index *L* = *a*^⊺^*Z* + *U* with *Y* | *L* ∼ Bernoulli(*F* (*L*)) for a strictly increasing link *F* : ℝ → (0, 1), the Bernoulli draw independent of everything else given *L*; detector *S* = *U* + *b*^⊺^*Z* + *ε*. A linear-index nuisance model stratifies on *T* = *w*^⊺^*Z*. Write Σ_*w*_ = Σ − Σ*ww*^⊺^Σ*/*(*w*^⊺^Σ*w*) for the covariance of *Z* left uncontrolled inside a stratum, and

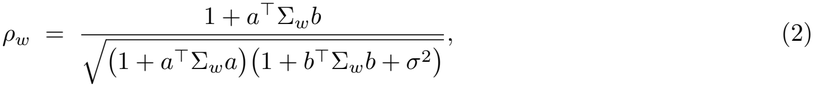

the correlation, inside a stratum, between the part of the label index and the part of the score that the strata leave uncontrolled.

#### Theorem 1

(Correlation law) *Conditional on T* = *t, the label index and the score are bivariate normal with correlation ρ*_*w*_, *the same for every t. Within a stratum the AUC of S is A*(*ρ*_*w*_; *m, v*), *where m and v are the stratum’s mean and variance of the label index; for each fixed* (*m, v*), *A is strictly increasing in ρ*_*w*_ *with* 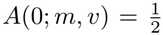. *Hence, for the pair-weighted estimand and for any nonnegative stratum weights with finite positive total mass*, 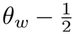 *has the sign of ρ*_*w*_. *With the target loading normalized to one, the estimate is below* 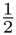 *if and only if a*^⊺^Σ_*w*_*b <* −1.

Appendix A gives the proof. In Family I the stratifier that omits 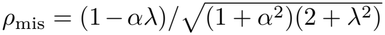, which gives the boundary *αλ* = 1. The sign is exact; the size is not a function of *ρ*_*w*_ alone, because *A* also depends on how much label variation the strata leave uncontrolled (*v* = 1 + *a*^⊺^Σ_*w*_*a*) and on the stratum mean. Two cells of Table 1 both have 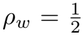 (the Family I truth at (1, 1) and the wrong model at (1, 0)) and read 0.619 and 0.648. We get the size by numerically integrating the analytic within-stratum AUC (Gauss–Hermite quadrature, no simulation) and check it against simulation.

### Verification

Table 1 and Figure 1 compare the quadrature against two simulation families with Gaussian data-generating variables that were run before the formula and its reversal boundary were derived. This is a retrospective comparison with earlier simulations, not an independent prospective validation of the theorem. The largest deviation over the 36 cells is 0.008 (0.005 in Family I). Every predicted reversal occurs, and the two *αλ* = 1 cells sit at 0.498 and 0.500 against a predicted 0.500. The worst cell, an estimate of 0.293 against a truth of 0.577, is (2, 2), predicted at 0.296. Family III’s law columns use the ideal population index of the six seen columns, while its simulation columns fit boosted trees in twenty bins, so that comparison mixes model and estimation error. New runs separate the two and extend the grid towards the boundary (Appendix B).

**Figure 1:**
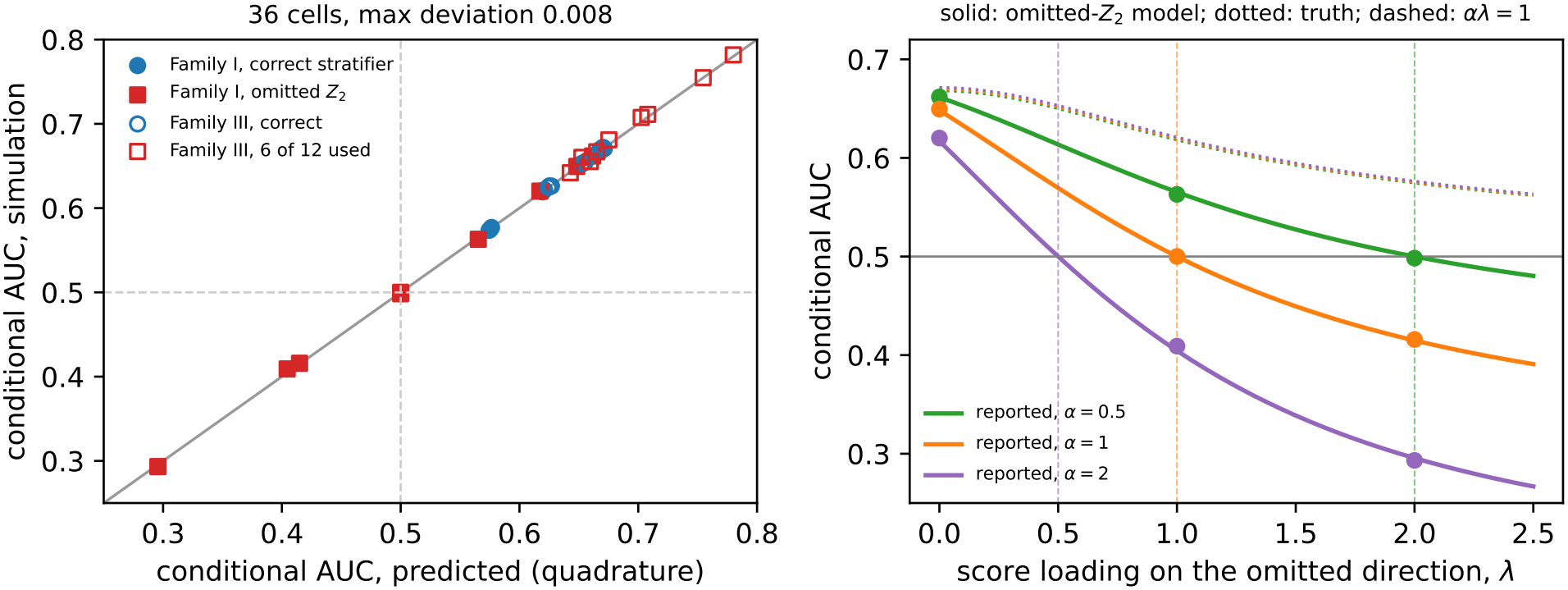
**Left:** the formula’s prediction (horizontal) against the simulated mean (vertical) for the 36 cells of Table 1; all lie within 0.008 of the diagonal. **Right:** the two-property example. Solid curves: the estimate of the model that omits *Z*_2_ as *λ* grows, one per *α*; dots: simulation means; dotted: the truth under the correct model; dashed verticals: the reversal boundary *αλ* = 1.

### Beyond the Gaussian model

The sign argument needs less than Gaussianity: within each stratum, a continuous label index, a strictly increasing link, and a score equal to *c* times the index plus noise independent of index and label (Proposition 4). The Gaussian model guarantees this and supplies (2); tree strata need not. For practical models we check the direction of the error empirically. In Family II (quadratic label index, linear model; §4) every condition mean with *λ >* 0 errs upward and every *λ* = 0 condition mean downward. On the real descriptors of five corpora, with synthetic labels and target (§4), we give the detector a loading on the part of the label’s direction that 16 features cannot absorb: *λ* ∈ {1, 2, 3} against the label, {1, 2} with it, or none. This is an adversarial stress test, not an estimate of how often reversal happens in practice. Logistic regression on 16 features and boosted trees on 62 report below the truth on every corpus with no loading and above it with the label’s sign (Table 2). Against the label at *λ* = 3 the 16-feature model falls below one half in 14 of 50 replicates (AFB: 9 of 10, mean 0.481 against a truth of 0.566), and at *λ* = 2 in 5 of 50. The direction agrees with the prediction in 590 of 600 replicate cells and in all 60 condition means, and 594 cells pass both checks. These are counts of experimental cells, not independent repetitions: detector settings share generated labels and nuisance fits within each replicate. These are deterministic reruns; historical counts differ slightly (Appendix H). Building the reference from 50 or 200 instead of 100 oracle bins leaves all 60 condition-mean directions and both reversal counts unchanged (replicate agreement 583–591 of 600; Appendix C).

**Table 2:** Reversal and inflation with practical nuisance models on real descriptors from five corpora and synthetic labels and target (§4; deterministic reruns). The first row gives oracle-stratum truth; other entries are mean signed errors (estimate minus fold-matched truth) over ten replicates. The detector loads against (*λ* = 3) or with (*λ* = 2) the label’s direction outside the 16 features’ span. Brackets count replicates with an estimate below one half and truth above it; counts are shown only when nonzero.

| nuisance model | detector loading | AFB | Kaggle | Wellgen | malaria | PCam |
|---|---|---|---|---|---|---|
| truth (oracle strata), AUC | against, λ = 3 | 0.566 | 0.600 | 0.647 | 0.599 | 0.578 |
| logistic, 16 features | against, λ = 3 | −0.085 [9/10] | −0.100 [5/10] | −0.093 | −0.063 | −0.058 |
|  | with, λ = 2 | +0.038 | +0.028 | +0.023 | +0.021 | +0.022 |
| boosted trees, 62 features | against, λ = 3 | −0.016 | −0.060 | −0.086 | −0.016 | −0.013 |
|  | with, λ = 2 | +0.038 | +0.035 | +0.026 | +0.029 | +0.033 |

## 4 What self-derived checks can establish

We first consider diagnostics restricted to the nuisance model’s inputs and outputs: balance on used covariates, propensity discrimination, and the self-stratified control. Two elementary facts show that exact balance and exact self-stratification can conceal an omitted covariate (Appendix A). Both are exact for a model that is correct on the covariates it uses and for exact level sets; with quantile bins they hold only approximately, and the experiments below measure how far.

First, a nuisance model that is correct for the covariates it uses is a balancing score for those covariates (Rosenbaum & Rubin, 1983). Within its strata the used covariates have the same distribution among positives and negatives, whether or not a covariate the model never saw drives the label (Proposition 6). Every balance statistic on the used covariates therefore sits at its null value while the adjusted AUC carries the error of §3. A wrong functional form on the used covariates is different: if the true propensity on those covariates is no longer a function of the fitted one, some function of the covariates ends up imbalanced, and a more flexible refit on the same covariates can detect that.

Second, a score stratified on itself is constant within every stratum, so its within-stratum AUC is one half by definition, whether the score is correct, misspecified, or arbitrary (Lemma 5). That is all the self-stratified control can ever report, in exact level sets. With finite bins it can depart from one half because of within-bin label gradients or limited overlap, so a pass is likely rather than guaranteed, and a coarse-bin failure is not evidence of omission: in a new Gaussian grid with 4 bins the checks reject 86% of correct-model replicates and 69% of omitted-input ones, while with 20 bins all 62 materially wrong replicates pass (Appendix B).

Table 3 collects the facts. A passed self-derived check tells you about overlap and, at most, about functional form on the used covariates. It cannot establish that omitted label drivers are absent.

**Table 3:**
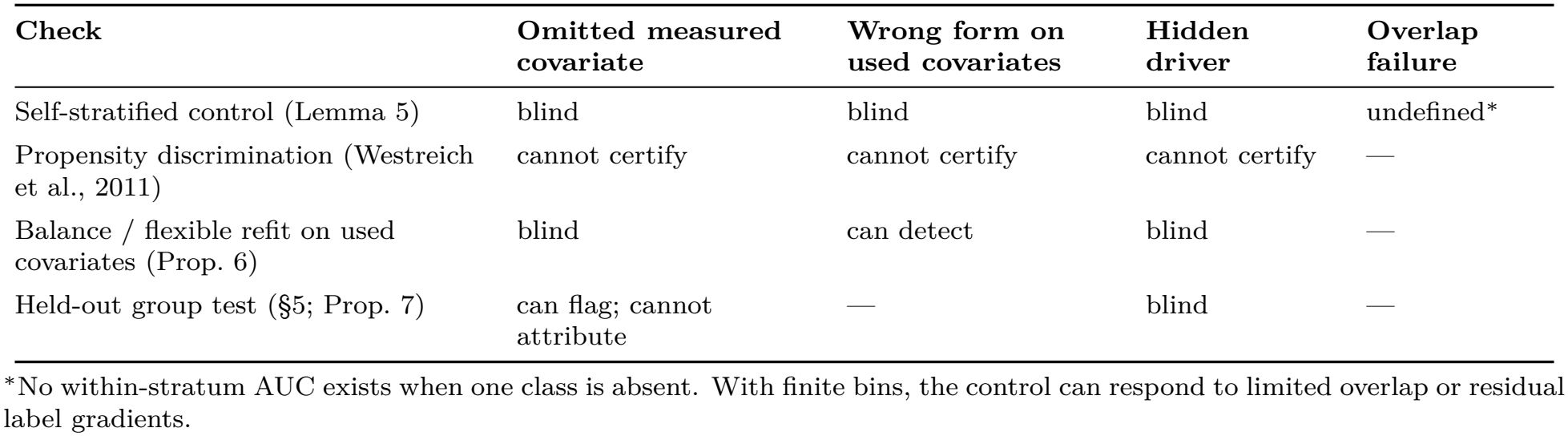
Scope of each diagnostic at the population level with exact strata. The omitted-covariate balance result assumes a correct propensity on the used covariates; the wrong-form column instead considers a fitted score that loses information in that propensity. “Blind” means that the check retains its null distribution. A hidden driver has no measured proxy; a correlated measured group may retain detectable signal (Proposition 7). Finite-bin behaviour is empirical.

| **Check** | **Omitted measured covariate** | **Wrong form on used covariates** | **Hidden driver** | **Overlap failure** |
|---|---|---|---|---|
| Self-stratified control (Lemma 5) | blind | blind | blind | undefined<sup>*</sup> |
| Propensity discrimination (Westreich et al., 2011) | cannot certify | cannot certify | cannot certify | — |
| Balance / flexible refit on used covariates (Prop. 6) | blind | can detect | blind | — |
| Held-out group test (§5; Prop. 7) | can flag; cannot attribute | — | blind | — |

<sup>\*</sup>No within-stratum AUC exists when one class is absent. With finite bins, the control can respond to limited overlap or residual label gradients.

### Ground-truth experiments

Three simulation families measure how often the checks miss a material error with 20 bins; the counts confirm, rather than add to, the exact results above. Family I is the two-covariate model of §3 (*α* ∈ {0.5, 1, 2}, *λ* ∈ {0, 1, 2}, ten replicates). The correct-input logistic model recovers the known *θ*_*B*_ within 0.02 in 93% of cells. The model that omits *Z*_2_ errs by up to 0.2998, yet **65 of 66** material cells pass both checks; the exception’s control reads 0.556, close to the correct-input maximum of 0.551. Family II (quadratic label index, linear model; worst error 0.139) and Family III (twelve AR(0.6) columns, six seen; 0.164) pass in all **35** and **40** material cells, so 140*/*141 pass overall. A separate five-replicate Family I sweep finds every material cell passing at all eighteen cutoff settings we tried (Appendix C). The control does move when overlap fails (0.874 on 17 MedMNIST instances with descriptor AUC 0.98–0.99, at balanced prevalence with four C3 strata), yet can remain near one half when directions are omitted.

### On real descriptors

Because Gaussian variables do not reproduce real descriptor geometry, we repeat the experiment on each corpus’s actual descriptors, using the 62 measured color, focus, texture, and edge properties. Only the label mechanism and the target latent are synthetic: the label’s log-odds combine the descriptor’s strongest principal direction with the top-ten component that the 16 color and focus features explain least well (out-of-fold *R*^2^ 0.44–0.73) and a standard-normal target, so the oracle nuisance index is known. Reference AUCs are finite-sample, fold-matched estimates using 100 oracle-index bins, not exact population values.

Conditioning on the known nuisance index recovers the truth with mean error 0.005 and maximum 0.010. Estimating the propensity from all 62 real features gives mean absolute errors of 0.011–0.037 per corpus against the oracle reference, with corpus-mean estimates lower and the screen always passing. The practical 16-feature logistic model errs by 0.040–0.065 wherever the label mechanism uses information outside those features, producing 24 material cells, and **all 24 pass**. Both errors follow the empirical direction described in §3.

## 5 Sensitivity to nuisance-model choice on real images

We now use real labels on the five corpora of Appendix C. With real labels the truth is unknown, so we ask what the checks and the available diagnostics can and cannot establish. All real-data quantities refer to the class-capped samples (at most 4000 tiles per class), and because (1) weights strata by *π*(1 − *π*), they can depend on that sampling prevalence; they are not estimates for a clinical population with a different prevalence. The Kaggle and Wellgen labels are source-level labels on local crops whose annotation trail was not retained, so a positive crop need not contain bacilli (Appendix C), and the AFB corpus is private. The comparisons below therefore describe how the estimator behaves on these collections, not clinical performance.

### Estimation and uncertainty

Bootstrap ranges use the 2.5th–97.5th percentiles of 1000 paired draws of the available groups (Appendix C), holding the fitted models fixed. Their population coverage is not established. Filename groups and near-duplicate clusters do not identify independent patients or slides; one PCam component contains 61% of tiles. Grouped refits provide a sensitivity analysis at these available units, not a patient-level validation.

### The disagreement

Table 4 holds the 16 features fixed and changes only the learner. On AFB, Kaggle and PCam both models pass both checks on fold-averaged values (in every individual split on Kaggle; see the “splits” column of Table 4 for the others), and logistic regression reads 0.044–0.055 higher than boosted trees, with the sign holding in 5 of 5 paired splits. Restricted to the individual splits in which both models pass both checks (“joint” in Table 4), the gap is 0.057 on AFB (3 splits), 0.048 on Kaggle (5) and 0.049 on PCam (3). On Wellgen the gap is similar but the tree model fails the overlap screen (22% of strata hold one label); on malaria the gap is small and reversed. The intervals hold the fitted models fixed, so they measure test-sample variability only; refitting with resampling units kept apart across training and test sets leaves mean gaps of 0.054 on AFB and 0.034 on Kaggle (Appendix F). Two correct population propensities on the same features would agree. Fitted propensities, however, change the conditioning groups and pair weights as well as introducing estimation and binning error. The observed gaps establish model sensitivity, not either estimate’s distance from a known correct adjusted AUC.

**Table 4:** Sensitivity to nuisance-model choice with the same 16 features (zero-shot detector; means over five splits). “Raw” is unadjusted AUC; C1 is logistic regression and C2 is boosted trees. Brackets are descriptive 95% paired cluster-bootstrap ranges with fits held fixed. “Checks” applies to fold-averaged values; “splits” counts individual passes for C1/C2. “Joint” gives the gap on splits where both pass, with the number of splits in parentheses. “Grouped” uses refits with available resampling units kept apart. PCam’s grouped AUC is undefined because its test sets contain only one class. Gaps are computed before rounding.

| corpus | raw | C1 | C2 | C1−C2 [bootstrap range] | checks | splits | joint | grouped |
|---|---|---|---|---|---|---|---|---|
| AFB | 0.913 | 0.894 | 0.838 | +0.055 [0.043, 0.068] | pass/pass | 5/3 | +0.057 (3) | +0.054 |
| Kaggle | 0.782 | 0.753 | 0.705 | +0.048 [0.032, 0.060] | pass/pass | 5/5 | +0.048 (5) | +0.034 |
| PCam | 0.838 | 0.721 | 0.677 | +0.044 [0.033, 0.084] | pass/pass | 4/4 | +0.049 (3) | undefined |
| Wellgen | 0.690 | 0.575 | 0.528 | +0.046 [0.001, 0.093] | pass/fail | 5/1 | +0.065 (1) | +0.026 |
| malaria | 0.934 | 0.846 | 0.861 | −0.015 [−0.024, 0.002] | pass/pass | 3/1 | −0.012 (1) | −0.012 |

### Bin count

The disagreement persists across the tested numbers of strata, while the checks’ verdicts change. Recomputing both models with *q* ∈ {5, 10, 20, 40} equal-count bins (Table 10, Appendix F) leaves the gap at 0.050–0.058 on AFB, 0.043–0.050 on Kaggle, 0.040–0.046 on PCam and 0.045–0.048 on Wellgen, positive in every split at every *q*, and between −0.015 and −0.010 on malaria. Coarse bins push the control out of its window (with *q* = 5 no model passes in more than one AFB split), while fine bins empty strata of one class (with *q* = 40 the tree model on Wellgen leaves 30% of strata with one label). Within-bin pairs, the only ones the estimator uses, make up 0.30 to 0.83 times 1*/q* of all positive–negative pairs (corpus means), almost independently of *q*. The share is lower for the tree model, whose strata separate the classes more sharply.

The disagreement is not specific to the zero-shot detector. For logistic probes trained on the real labels over BiomedCLIP or DINOv2 (Oquab et al., 2024) embeddings, C1 minus C2 is 0.022–0.117 on AFB, Kaggle, PCam and Wellgen, with intervals excluding zero, and near zero on malaria; weaker zero-shot detectors give gaps of mixed sign. A different feature set is a different quantity: the same tree learner on 62 rather than 16 features moves the estimate by 0.051–0.073 on AFB, Kaggle and PCam, and with all 95 features C4 fails at least one check on four of five corpora (Appendices C and F).

### What the diagnostics show

The two checks we study are not the only diagnostics available, and broader ones do separate the models (Table 5). Within each of the 20 bins we compute the absolute standardized mean difference of every feature between positives and negatives, using the bin’s own pooled SD, and average over bins with pair weights. Small bins make this statistic noisy, so we compare it with a floor obtained by permuting labels within bins (20 permutations), which keeps bin sizes and class counts but removes any imbalance. C2’s imbalance sits close to the floor on every corpus (excess 0.01–0.05), while C1’s exceeds it by 0.09–0.19. C1’s calibration error is also 1.9–4.4 times C2’s on four corpora (PCam is the exception), and C2’s propensity still discriminates the label inside C1’s strata (AUC 0.76–0.94; the reverse, 0.49–0.70). These diagnostics all point to C1 fitting the label less well. They are warnings about fit, not proof that C2 gives the correct adjusted AUC: C2 is near the floor, not at it, and on malaria both cross-model directions retain signal (0.759 and 0.696). C2|C1 in Table 5 is the adjusted AUC of C2’s propensity used as a score within C1’s strata, and C1|C2 the reverse. Hosmer–Lemeshow tests reject both models on most corpora, but their nominal *p*-values ignore clustering.

**Table 5:** Fit diagnostics for the same-feature models (zero-shot, means over five splits). Slash-separated pairs, including floors, are ordered C1/C2. |SMD| is the pair-weighted mean over bins and features of the absolute within-bin standardized mean difference, with its label-permutation floor in brackets. ECE is expected calibration error. C2|C1 uses C2’s propensity as a score within C1’s strata; C1|C2 reverses these roles.

| corpus | \|SMD\| C1/C2 [floor] | ECE C1/C2 | C2\|C1 | C1\|C2 |
|---|---|---|---|---|
| AFB | 0.41/0.26 [0.23/0.25] | 0.076/0.040 | 0.859 | 0.536 |
| Kaggle | 0.29/0.23 [0.18/0.20] | 0.091/0.046 | 0.835 | 0.557 |
| PCam | 0.33/0.25 [0.20/0.21] | 0.037/0.039 | 0.797 | 0.529 |
| Wellgen | 0.28/0.25 [0.19/0.22] | 0.100/0.023 | 0.942 | 0.493 |
| malaria | 0.38/0.29 [0.25/0.23] | 0.073/0.021 | 0.759 | 0.696 |

**Table 6:** Corpora as used: AFB (Heinzman et al., 2026), Kaggle (Quinn et al., 2016), Wellgen (Lin & Wellgen Medical, 2024), malaria (Rajaraman et al., 2018), PCam (Veeling et al., 2018). Tiles are class-capped at 4,000 per class. “Groups” gives the resampling unit for cluster bootstraps; near-duplicate clusters are computed on DINOv2 embeddings (Oquab et al., 2024) at cosine threshold 0.95. None is a recorded slide or patient identifier.

| corpus | stain, target | tiles | positives (share) | public | groups |
|---|---|---|---|---|---|
| AFB | ZN, acid-fast bacilli | 6,074 | 2,074 (0.34) | no | 5,502 near-duplicate clusters |
| Kaggle | ZN, acid-fast bacilli | 8,000 | 4,000 (0.50) | yes | 344 filename-derived image groups |
| Wellgen | ZN, acid-fast bacilli | 8,000 | 4,000 (0.50) | yes | 354 fields |
| malaria | Giemsa, parasitized cells | 8,000 | 4,000 (0.50) | yes | 4,884 near-duplicate clusters |
| PCam | H&E, tumor | 8,000 | 4,000 (0.50) | yes | 2,939 near-duplicate clusters |

### A known shortcut on real images

To test a controlled shortcut on real images, we multiply the red channel by 1 + *δ* and the blue by 1 − *δ* (*δ* ∈ {0.10, 0.25}) on a random subset of tiles, shifting each positive with probability 0.8 and each negative with 0.2 (with the label), the reverse (against it), or both with 0.5 (independent), and score the zero-shot detector and a probe trained on the shifted images. The detector’s loading on the shift is the mean score change between paired shifted and original images, divided by the original-score SD. For the zero-shot detector it is measured on all tiles at each shift strength; for each supervised probe it is measured only on its held-out test fold, separately by condition. C1 and C2 fitted to original-image features cannot see the shift; each one’s reference crosses its propensity bins with the known shift indicator, a model-specific reference rather than a single correct AUC. Deviations, averaged over five splits, take the direction predicted from the detector’s loading on the shift in 78 of 80 label-dependent cells, including all 49 deviating by more than 0.02 (largest inflation 0.127), and 171 of the 246 fold evaluations that individually deviate by more than 0.02 pass both checks. With a label-independent shift deviations average 0.002; features of the shifted images lower the mean absolute deviation from 0.036 to 0.016. This tests the predicted direction on real images, not the Gaussian formula (Appendix F).

### Exploratory model selection

We also compare two members of each model class by cross-fitted label log-loss, selecting a class when its paired per-image difference has |*z*| ≥ 2 and abstaining otherwise (Appendix E). The rule uses only training labels and nuisance features. It selects trees in every split on all five corpora, including malaria, where trees give the higher adjusted AUC. On AFB the class-selected estimates are 0.890 (linear) and 0.842 (tree), a gap of 0.048 [0.036, 0.060]. Validation shows limited downstream benefit: mean adjusted-AUC error is 0.0314 against 0.0328 for always choosing trees over 480 replicates of a linear-favouring design, and the abstention calibration criterion fails (Table 7). We therefore report both class-selected estimates and use label fit as a diagnostic.

**Table 7:** Validation of the label-fit selection rule on each corpus’s real 16-feature geometry with synthetic labels. A condition is informative when the true error gap between classes is at least 0.01. The first two rows count decided conditions; the final row counts replicate decisions and conditions in which the rule beats trees. Errors are mean absolute adjusted-AUC errors over decided conditions (rule versus always choosing trees); for the linear-favouring design, over all 480 replicates with abstentions counted as trees. The frozen replication’s verdict counts come from its 12-replicate block and its errors from its 24-replicate block; the bracket is an exact binomial 95% interval.

| block | informative (lin./tree) | decided | correct | error: rule / trees |
|---|---|---|---|---|
| original (6 replicates) | 11 (2/9) | 7 | 7/7 [0.59, 1.00] | 0.0195 / 0.0218 |
| frozen replication (12; 24) | 10 | 7 | 6/7 | 0.0196 / 0.0198 |
| linear-favouring (12) | 4 (4/0) | 374 linear, 106 abstain | 4/4 beat trees | 0.0314 / 0.0328 |

### Held-out tests (exploratory)

A check can see an omitted measured group only by using it (Table 3). A split-family test that adds half of a held-out descriptor group to the propensity detects strong omitted signal in Gaussian designs, but cannot tell nuisance from target signal, and on real descriptors with nothing omitted it rejects on 16.9% of halvings, so we draw no omission conclusions from it (Appendix D).

### A checklist for evaluators

1. Choose the adjustment set with clinical input; flag properties that may carry disease signal, such as morphology, and report results with and without them.
2. Fit at least two classes of nuisance model on the same features.
3. Check overlap, but do not treat the self-stratified control as evidence that the model is right.
4. Score each model’s propensity inside the other’s strata as a fit check, allowing for residual signal from finite bins.
5. Report the raw AUC, each model’s adjusted AUC, and per-split check results. State the resampling unit and whether uncertainty includes refitting.
6. Calibrate held-out tests on relevant nulls, aggregating splits as in the evaluation; investigate flagged features before concluding omission.

This is a reporting framework, not a validated method for finding the correct adjusted AUC: the cluster intervals are descriptive, whether the adjustment set is right is a clinical judgment, and better label fit carries no guarantee of a more accurate estimate.

## 6 Limitations and conclusion

The theorem characterizes reversal under exact linear-index conditioning in the Gaussian model. Proposition 4 extends the sign argument under an independent-noise condition that practical tree strata need not satisfy. Inflation and attenuation relative to correct adjustment remain empirical findings in the tested designs.

The real-data results demonstrate sensitivity to model choice, not a known error in either estimate. The colour-shift references are model-specific, label-fit selection provides only small improvements, and the held-out test is not calibrated for real descriptors. Available groups do not recover unrecorded patient or slide identities, and the bootstrap ranges exclude refitting uncertainty. The corpora are limited to microscopy and pathology tiles. In addition, texture and morphology can carry disease signal; adjusting for them may remove diagnostically relevant information as well as shortcuts (Schisterman et al., 2009).

Propensity-adjusted AUC should therefore be reported with the adjustment set, raw AUC, alternative nuisance models, and the scope of the diagnostic checks. Exact balance on used covariates cannot certify completeness of the adjustment set; broader fit diagnostics can still reveal problems with how those covariates are modeled.

## Data and code availability

We use local preprocessed versions of five pathology tile corpora (provenance limitations in Appendix C). Public source collections include the Ziehl–Neelsen smear photographs of Quinn et al. (2016) in their Kaggle distribution, the Wellgen Medical TB smear image set (Lin & Wellgen Medical, 2024), PatchCamelyon (Veeling et al., 2018), and the NIH malaria thin-smear set (Rajaraman et al., 2018); MedMNIST v2 (Yang et al., 2023) is used for one overlap example. The fifth, the Ziehl–Neelsen-stained acid-fast bacilli corpus (AFB) described by Heinzman et al. (2026), is not publicly available. The Gaussian simulations need no image data; semi-synthetic experiments require the corresponding cached image descriptors. The analysis package nuisauc, run instructions, and aggregate reference tables are staged for release at https://github.com/calebheinzman/nuisance-auc. The Gaussian simulations and quadrature can be rerun from code alone. Readers can check reported summaries against the aggregate reference tables with the supplied analysis code. Repeating the image experiments requires separately obtained inputs. The public release excludes per-tile inputs and restricted AFB images. Our local Kaggle and Wellgen crops cannot be reconstructed exactly from the public sources because the source versions and crop-level annotation trail were not retained (Appendix C).

## Ethics statement

This work uses previously collected microscopy images and involves no interaction with human subjects. The AFB images were collected and de-identified under protocol 29897, Teaching computer to recognize diagnostic features of whole slide images, approved as exempt (category 4) by the Saint Louis University Institutional Review Board on March 27, 2019 (assurance FWA00005304). The other corpora are public releases.

## A Exact statements and proofs

### Model

*Z* ∼ *N* (0, Σ) in ℝ^*p*^ with Σ positive definite, a target latent *U* ∼ *N* (0, 1), and score noise *ε* ∼ *N* (0, *σ*^2^) with *σ*^2^ *>* 0, mutually independent. The label index is *L* = *a*^⊺^*Z* + *U* with *a* ≠ 0 and *Y* | *L* ∼ Bernoulli(*F* (*L*)) for a strictly increasing link *F* : ℝ → (0, 1) (logistic in all our experiments), the Bernoulli draw independent of everything else given *L*. The detector is *S* = *U* + *b*^⊺^*Z* + *ε*. A linear-index nuisance model stratifies on *T* = *w*^⊺^*Z* with *w* ≠ 0; when it is correctly specified on a subset *Z*_used_ of the columns, its population index direction is that of *E*[*a*^⊺^*Z* | *Z*_used_] (Brillinger, 1983; Li & Duan, 1989). Σ_*w*_ = Σ − Σ*ww*^⊺^Σ*/*(*w*^⊺^Σ*w*) is the covariance of *Z* given *T, ρ*_*w*_ is (2), and |*ρ*_*w*_| *<* 1 because *σ*^2^ *>* 0.

#### Theorem 1

**(full statement)** *Conditional on T* = *t, the centered label index and score are bivariate normal with correlation ρ*_*w*_, *which does not depend on t. Let m*_*w*_(*t*) = *E*[*L*| *T* = *t*] *and v*_*w*_ = Var(*L* | *T* ) = 1 + *a*^⊺^Σ_*w*_*a. The within-stratum AUC of S is A*(*ρ*_*w*_; *m*_*w*_(*t*), *v*_*w*_), *where for each fixed* (*m, v*) *the function ρ* ↦ *A*(*ρ*; *m, v*) *is strictly increasing on* (−1, 1) *with* 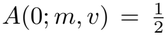. *Consequently, for the pair-weighted estimand and for any nonnegative stratum weights with finite positive total mass*, 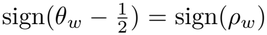. *If w ∝ a, then a*^⊺^Σ_*w*_*a* = *a*^⊺^Σ_*w*_*b* = 0 *and ρ*_*a*_ = (1 + *b*^⊺^Σ_*a*_*b* + *σ*^2^)^−1*/*2^ *>* 0.

*Proof. Step 1: within-stratum moments*. Given *T* = *t*, the residual 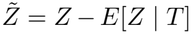 has covariance Σ_*w*_ and does not depend on *t*. The centered label index and score are

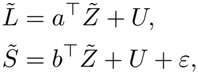

with variances and covariance

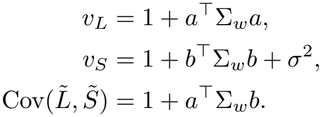

Their correlation is therefore *ρ*_*w*_ of (2), the same in every stratum.

*Step 2: decomposition*. Because 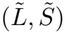 is bivariate normal,

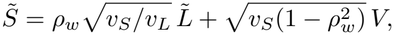

with *V* ∼ *N* (0, 1). *V* is independent of 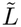, and of *Y*, which depends on the stratum’s variables only through *L*.

*Step 3: AUC as an expectation*. Within the stratum, positives and negatives have class-conditional index densities

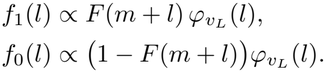

For independent draws 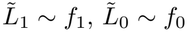 and 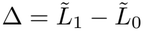, Step 2 gives

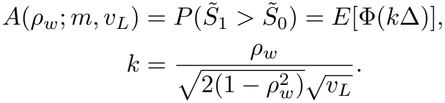

At *ρ*_*w*_ = 0, *k* = 0 and 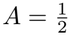.

*Step 4: monotonicity*. Differentiating,

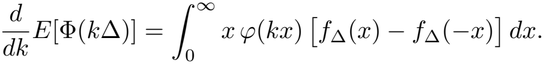

With *g* = *f*_1_*/f*_0_ *∝ F* (*m* + *l*)*/*(1 − *F* (*m* + *l*)), which is strictly increasing,

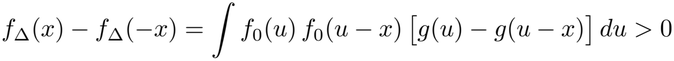

for *x >* 0. So *E*[Φ(*k*Δ)] is strictly increasing in *k*, and *k* is strictly increasing in *ρ*_*w*_ with the same sign.

*Step 5: pooling*. Every stratum’s AUC minus one half therefore has the sign of *ρ*_*w*_, which is common to all strata, and any nonnegatively weighted average with positive total mass inherits it. If *w* ∝ *a* then Σ_*w*_*a* = 0, which gives the last claim.

#### Corollary 2

(Regimes of the correlation) *Let a*^⊺^Σ_*w*_*a >* 0 *and normalize the detector’s target loading to one. (i) If b* = 0, *then ρ* = ((1 + *a*^⊺^Σ_*w*_ *a*)(1 + *σ*^2^)^−1/2^ *< ρ* = (1 + *σ*^2^)^−1/2^. *(ii) The estimate is below* 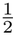 *if and only if a*^⊺^Σ_*w*_*b <* −1, *and equals* 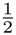 *when a*^⊺^ Σ _*w*_*b =* − 1.

*Proof*. (i) With *b* = 0 the numerator of (2) is 1 and the first denominator factor exceeds 1; *ρ*_*a*_ is the last claim of Theorem 1 with *b* = 0. (ii) follows from the sign of the numerator and Theorem 1.

*Remark* 3 (Observed, not proven). Part (i) compares correlations, not adjusted AUCs: the wrong stratifier also raises the uncontrolled label variance *v*_*L*_ and changes the stratum means, on which *A* depends, so attenuation of the pooled estimate below the truth is an observation (every *b* = 0 cell of Table 1 and every no-loading condition mean of the semi-synthetic grid), not a theorem. When *a*^⊺^Σ_*w*_*b >* 0 the shortcut is credited and the estimate can exceed the truth, as Family III shows at *λ >* 0; this too is exhibited, not an ordering theorem for two stratifiers.

For Family I (Σ = *I, a* = *α*(1, 1), *b* = *λ*(1, −1), *σ*^2^ = 1) the correct stratifier *T* = *Z*_1_+*Z*_2_ has 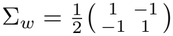 giving *ρ* = (2 + 2 *λ* ^2^)^−1/2^; the stratifier that omits *Z*_2_ Σ_*w*_ = diag(0 1), giving *a*^⊺^Σ_*w*_*b* − *αλ* and 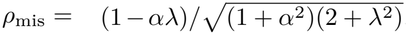. The covariance 1 + *a*^⊺^Σ_*w*_*b* is the omitted-variable term of linear regression and the reversal a Simpson-type sign change under partial adjustment, distinct from bias amplification (Pearl, 2010; Ding et al., 2017).

#### Proposition 4

(Sign without Gaussianity) *Fix a stratum with label-index mean m. Let the centered label index* 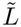 *have a density, let Y* | 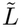 ∼ Bernoulli(*F* (*m* + 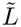)) *with F strictly increasing, and suppose the score is* 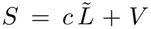, *where V has a density and is independent of* (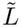, *Y* ). *Then* 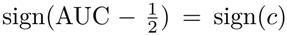, *which equals* sign Cov(*S*, 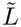) *when* 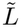 and *V have finite variance. For nonnegative stratum weights with finite positive total mass, the weighted average of stratum AUCs minus one half has the sign shared by the strata’s coefficients c*.

*Proof*. Draw (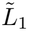, *V*_1_) from the positive class and (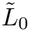, *V*_0_) from the negative class, independently, and write 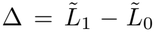 and *W* = *V*_1_ − *V*_0_. *W* is independent of Δ, continuous and symmetric about zero, so its distribution function *G* satisfies *G*(*x*) + *G*(−*x*) = 1, and

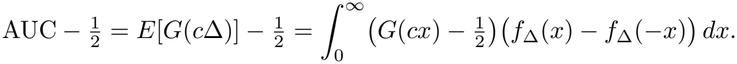

The argument of Step 4 of the proof of Theorem 1 gives *f*_Δ_(*x*) − *f*_Δ_(− *x*) ≥ 0 for *x >* 0 without Gaussianity of the index, and the inequality is strict on a set of positive measure near zero, because by continuity of translation in *L*^1^ the density *f*_0_ and its translate by a small *x* overlap on a set of positive measure. 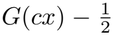 has the sign of *c* for *x >* 0 and is nonzero for every *x >* 0 when *c* ≠ 0: truncating the density of *V* to a bounded set of positive mass, the autocorrelation of the truncation is continuous and positive at zero, so *P* (0 *< W < t*) *>* 0 for every *t >* 0. Hence the integral has the sign of *c*; with finite variances, 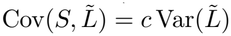.

Joint Gaussianity with linear-index strata guarantees this structure with one *c* for all strata, and supplies the closed form (2) and the boundary *a*^⊺^Σ_*w*_*b* = −1. With non-Gaussian descriptors the residual of the score on the index is uncorrelated with it but need not be independent, and with tree strata *c* can vary across strata. Lemma 5 and the balance statement of Proposition 6 need no Gaussianity.

#### Lemma 5

(Self-stratification) *Let R be any real-valued measurable function of X and stratify on G* = *R*(*X*). *For every stratum with* 0 *< P* (*Y* =1 | *G*=*g*) *<* 1, *the within-stratum AUC of R as a score for Y, with ties given half credit, equals exactly* 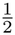; *hence any nonnegatively weighted stratum average equals* 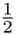, *for R correct, misspecified, or arbitrary*.

*Proof*. Given *G* = *g, R* = *g* almost surely, so every positive–negative pair is tied and the tie-corrected AUC is 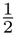. Finite bins only approach this (in the smooth Gaussian example *q* = 10, 20, 100 give 0.528, 0.514, 0.502), and departures reflect within-bin label gradients, sampling noise, or limited overlap.

#### Proposition 6

(Self-derived checks) *Partition the descriptor as Z* = (*Z*_*A*_, *Z*_*B*_) *and let the nuisance model use Z*_*A*_ *only. Suppose it is correct given what it uses, ê* = *e*_*A*_ := *P* (*Y* =1 | *Z*_*A*_). *Then Y* ⊥ *Z*_*A*_ | *e*_*A*_, *so the null hypothesis of every within-stratum balance test on functions of Z*_*A*_ *holds exactly; the self-stratified control equals* 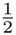 *on every stratum with* 0 *< e*_*A*_ *<* 1 *(and in aggregate when such strata carry positive weight); and this holds identically whether or not Z*_*B*_ *enters the true label mechanism. In the Gaussian model of Theorem 1, e*_*A*_ *is a strictly increasing function of the linear index g*^⊺^*Z*_*A*_, *with g as in Appendix B for Z*_*S*_ = *Z*_*A*_, *so whenever g* ≠ 0 *the adjusted AUC has the correlation and sign relative to one half characterized by Theorem 1 with w* = (*g*, 0). *If instead e*_*A*_ *is not a function of ê (a wrong functional form on Z*_*A*_ *that loses information in e*_*A*_*), then* 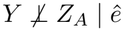, *and a balance statistic on some function of Z*_*A*_, *or a more flexible refit on Z*_*A*_, *departs from its null value*.

*Proof*. Since *e*_*A*_ is a function of *Z*_*A*_, *P* (*Y* =1 | *Z*_*A*_, *e*_*A*_) = *e*_*A*_ = *P* (*Y* =1 | *e*_*A*_), so *Y* ⊥ *Z*_*A*_ | *e*_*A*_ (Rosenbaum & Rubin, 1983), which is the null of every balance statistic on *Z*_*A*_, and Lemma 5 gives the control; *Z*_*B*_ never enters, and Theorem 1 applies to the index of *e*_*A*_. Conversely *P* (*Y* =1 | *Z*_*A*_, *ê*) = *e*_*A*_, so *Y* ⊥ *Z*_*A*_ | *ê* if and only if *e*_*A*_ is a function of *ê*, and a form under which it is not leaves *e*_*A*_ itself, which a flexible refit estimates, imbalanced.

## B Families II and III

Family III draws *Z* ∈ ℝ^12^ with AR(1) covariance Σ_*ij*_ = 0.6^|*i*−*j*|^ and *U, ε* ∼ *N* (0, 1), all independent. The label’s log-odds are *α a*^⊺^*Z* + *U* with *a* ∝ **1** scaled so that *a*^⊺^*Z* has unit variance, and the detector is *U* + *λ h*^⊺^*Z* + *ε* (so *b* = *λh*) with *h* ∝ (0, …, 0, 1, …, 1), the mean of the six hidden columns scaled to unit variance; *α* ∈ *{*0.5, 1, 2}, *λ* ∈ *{*0, 1, 2}, *n* = 20,000 per replicate and ten replicates, with 200-iteration boosted trees wherever trees are fit; Families I and II use the same sample size and learners. The correct model sees all twelve columns; the archived misspecified model is boosted trees on the six seen columns *Z*_*S*_, while the quadrature evaluates the ideal population balancing index on that subset, so the comparison includes tree-estimation and finite-bin effects. That index is *m* = *E*[*a*^⊺^*Z* | *Z*_*S*_] = *g*^⊺^*Z*_*S*_ with 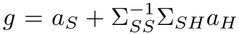 (Brillinger, 1983; Li & Duan, 1989), and (2) is evaluated with label loading *a*_law_ = *αa* detector loading b = *λh, w* = (*g*, 0), and 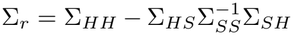. Thus 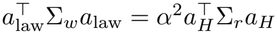 and 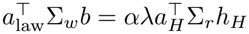; the oracle stratifier uses *w* = *a*, and the seen block explains a fraction 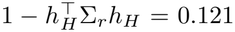 of the hidden mean’s variance. At *λ* = 0 the law predicts attenuation of 0.002, 0.009, 0.028 against 0.002, 0.013, − 0.030 observed, and at (2, 2) inflation of +0.153 against +0.156 observed (Family III columns of Table 1); the quadrature and an independent Monte Carlo check of the law columns are in the code release.

Family II draws *Z*_1_, *Z*_2_, *U, ε* ∼ *N* (0, 1), sets the label core to 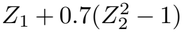 standardized to unit variance, the label’s log-odds to *α ·* core + *U*, and the detector to 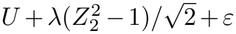, on the same grid and replicate count as Family III. The correct model is boosted trees on (*Z*_1_, *Z*_2_); the wrong model is logistic regression on (*Z*_1_, *Z*_2_), which cannot represent 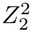. No exact formula applies; the sign prediction uses the detector’s loading on the missed functional 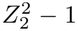, which for *λ >* 0 has the label’s sign (inflation) and for *λ* = 0 is absent (attenuation).

### Denser grid, estimation error, and bins

A new Family I grid of 36 settings, with *αλ* from 0 to 4 and several values close to the boundary, matches the formula for the stratifier that omits *Z*_2_ to the mean of ten replicates within 0.0068, and the correct-stratifier truth within 0.0099; near *αλ* = 1 individual replicates fall on either side of one half. Over 15 Family III settings, refitting the wrong model as logistic regression on the six seen columns, which estimates the ideal index, gives a largest formula–simulation discrepancy of 0.0063, against 0.0071 with boosted trees. On bins: with *q* = 4 the overlap screen and control pass only 14% of correct-model replicates and 31% of omitted-input replicates, while with *q* = 20 and *q* = 50 every replicate passes, including 62 of 62 and 63 of 63 materially wrong ones.

## C Corpora, descriptors, and protocol details

### Local data provenance

Experiments use cached descriptors and the labels supplied with local preprocessed tiles, not re-adjudicated for this study; the Kaggle and Wellgen names identify local derivatives of the cited collections whose source-version and crop-level annotation trail was not retained, so source-level positivity need not establish that every crop contains bacilli.

### Descriptors

The 16 color and focus features are the tile mean and standard deviation of seven per-pixel channels (red, green, blue, HSV saturation and value, two stain-unmixing channels) plus the variance of the Laplacian and the gray standard deviation (focus); the 62-feature descriptor (C3; group *A* of the held-out tests) adds 46 texture and edge statistics. C4 adds 18 frequency features (radial Fourier band energies per RGB channel plus anisotropy) and 15 object-level morphology summaries; each of these two sets serves as a held-out group *B*.

### Strata and folds

Quantile strata number 20 when the test fold holds at least 1,000 tiles, 10 when at least 200, and 4 otherwise (every real-corpus test fold uses 20). Real-corpus values are means over five repeated stratified 75/25 train/test splits (seeds 0–4), whose test sets can overlap, so between-split variation is not an independent-sample standard error; the nuisance model is fit on training tiles, strata are cut on test tiles, and inner four-fold cross-fitting is disjoint within each training split.

### Semi-synthetic grid

For each corpus and replicate the label’s log-odds are the standardized sum of the descriptor’s first principal direction *a*_1_ and the component *a*_*lo*_ among PCs 2–10 least predictable from the 16 color and focus features, plus a standard-normal target latent *u*. The detector is *u* + *ε, u* + *λ*(*a*_1_ − *a*_*lo*_) + *ε* with *λ* ∈ *{*1, 2, 3} (against the label), or *u* + *λ*(*a*_1_ + *a*_*lo*_) + *ε* with *λ* ∈ *{*1, 2} (with it); the oracle index, logistic regression on the 16 features, and boosted trees on the 62 are compared with the fold-matched oracle truth over ten replicates (600 non-oracle cells; Table 2). The archived generators seeded with Python’s process-dependent hash(corpus), so their draws cannot be reconstructed; all three-regime grid counts we report come from deterministic reruns with a SHA-256 corpus seed, and Appendix H documents the historical counts. The ten cells whose error sign disagrees with the rule are single replicates: one in each model’s no-loading condition on Wellgen (condition means −0.007 and −0.0065) and eight in tree-model conditions on malaria, PCam and AFB, where the 62 features absorb the loaded direction almost completely (condition means −0.005 to −0.017). The six cells failing the checks are one Kaggle tree-model replicate whose control reads 0.556. With the fold-matched truth computed from 50 or 200 oracle bins instead of 100 (same draws), truths move by at most 0.017 (mean 0.004); replicate direction agreement is 591 and 583 of 600, every condition mean keeps its predicted sign, and the reversal counts stay 14/50 and 5/50.

### Self-training

The self-trained score pseudo-labels the top and bottom 20% of tiles by the prompt score and trains a logistic student twice on the same frozen embeddings without ground-truth labels, using the whole corpus (test-tile embeddings included) before the evaluation splits: transductive self-training, not an inductive held-out-patient evaluation. Refitting the student on training-fold pseudo-labels only changes each corpus’s mean raw, C1 and C2 AUCs by less than 0.003.

### Overlap-screen convention and enrichment

Enrichment adds held-out groups to the nuisance model; C4 implements it and fails at least one check on four of five corpora (§5): malaria, AFB, and Wellgen because 0.22, 0.25, and 0.59 of strata contain one label only against the 0.20 threshold, and Kaggle, with adequate overlap, because its control reads 0.551, just outside the window. At a 30% threshold AFB would pass both checks and malaria, whose control reads 0.557, would still fail, so C4’s pass count depends on the conventions of §2. On PCam, where C4 passes, it moves the estimate by at most 0.026 relative to C3, against spreads of up to 0.112 among models passing both checks on the same corpus. In a Family I sensitivity sweep (5 replicates) at stratum multipliers *m* ∈ *{*0.5, 1, 2}, misspecified cells with error above 0.05 number 29–30 per multiplier, and at every combination with degenerate-strata thresholds {0.10, 0.20, 0.30} and control windows [0.45, 0.55], [0.40, 0.60] the fraction of material-error cells passing the screen is 1.00 (18 of 18 settings).

## D Held-out tests: statistic, calibration, and Proposition 7

### Statistic

Held-out balance (HOB) fits the propensity on group *A* (plus, for the split-family test, half of group *B*), cuts strata on the test fold, and measures the within-stratum label signal of a feature set *B*^*′*^ as 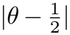, where *θ* is the adjusted AUC, within those strata, of a cross-fitted logistic probe on *B*^*′*^ (4-fold, never scored on tiles it was fit on); EXC is the maximum over the two half assignments of HOB(excluded half) − HOB(included half). The threshold 0.0167 is the 95th percentile of EXC over 180 correct-model calibration cells (Gaussian specificity, detection-limit, and split-stability studies); the semi-synthetic studies recalibrated on their own oracle cells to 0.01947 and 0.01760 (sensitivity study). On the six-column generative model of the sensitivity study, HOB detects 100% of omissions represented in the held-out group (90 of 90) at a pooled 5% false-positive rate (9 of 180); on the real corpora it rejects for groups that were included in the propensity in 7 of 10 corpus–group cases, the floor that the split-family test subtracts.

### Calibration and limit of detection

Against ground truth the split-family test detects 100% of diffuse and concentrated omissions, with a 7.2% false-positive rate pooled over the sensitivity study’s correct-model and functional-form cells (13 of 180; 6.7% on correct-model cells alone). In the separate specificity design (a correct propensity on *A* = (*Z*_1_, …, *Z*_4_) with logit *P* (*Y* =1) = *α*(*Z*_1_ + *Z*_2_) + *U* and a held-out group of four columns carrying *ρU* in *k* ∈ *{*1, 2, 4} of them, *ρ* ∈ *{*0.3, 0.6, 1.0, 1.5}, *α* ∈ *{*0.5, 1, 2}, ten replicates, 390 cells, held-out-probe AUCs 0.55–0.67 by regime) the rejection rates at the archived threshold are noise 0.067, diffuse 1.00, and concentrated 1.00 for *k* = 1 and *k* = 2, and the EXC statistic itself rises with *ρ* in concentrated regimes (Spearman +0.96 at *k*=1) and falls in the diffuse regime (− 0.60), qualitatively resembling the latent-covariance behaviour of the proposition below; these EXC trends are empirical. Detection rises steeply with driver strength (Spearman *ρ* = 0.927), reaches 80% at raw driver AUC 0.568, and across a 240-cell sweep no non-rejecting cell has true bias above 0.025 (95th percentile 0.019): empirical calibration values, not bounds. The semi-synthetic results below use the separately calibrated 0.01947 threshold and establish no transferable error bound for non-rejecting cells. Along the frequency group’s main axis *b*_1_ (*R*^2^ given the descriptor 0.67– 0.97) the nuisance model absorbs most of the driver on four of five corpora (mean bias 0.009–0.019); malaria, at *R*^2^ = 0.67, reaches 0.031 with the split-family test rejecting 5% of the time, missing the criterion of error at most 0.03 or detection at least 80%, as in the new confirmation (Appendix H). Along the residual 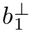 (out-of-fold *R*^2^ predicting the residual from held-out group *B*, 0.05–0.41) the rejection rate stays at most 0.06 at every strength while the full driver biases the estimate by up to **0.071** (non-rejecting-cell 95th percentile 0.056; other directions in Appendix H).

#### Proposition 7

(Target-proxy covariance) *Let U* ∼ *N* (0, 1) *be the target latent, let the held-out group carry only target proxies, B*_*j*_ = *ρU* + *η*_*j*_ *with η*_*j*_ *i*.*i*.*d. N* (0, 1) *independent of U and of the nuisance features A, with U* ⊥ *A, and let the latent outcome be Y* ^*^ = *h*(*A*) + *U* + *ε with Y* = **1 {***Y* ^*^ *>* 0}, *where ε has finite variance and is independent of* (*A, U, η*), *and h*(*A*) *has finite variance. There is no omitted nuisance in this population model. Split B into halves and stratify exactly on the population index m* = *h*(*A*) + *E*[*U* | *B*_inc_]. *Then, conditionally on m: (i)* Cov(*B*_inc_, *Y* ^*^ | *m*) = 0 *exactly; (ii) each excluded column retains conditional covariance* Cov(*B*_exc,*j*_, *Y* ^*^ | *m*) = *ρ* Var(*U* | *B*_inc_), *which has the sign of ρ and is positive for the ρ >* 0 *of every design here. Writing k* = *k*_inc_, *the excluded-minus-included covariance equals c*(*ρ*) = *ρ/*(1 + *kρ*^2^): *on ρ* ≥ 0, *for k >* 0 *it increases until* 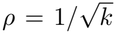 *and decreases thereafter; for k* = 0 *it increases linearly. Pure-noise columns contribute zero and are not counted in k*.

*Proof*. With 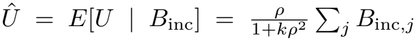 and *v* = (1 + *kρ*^2^)^−1^, the Gaussian residual *U* − *Û* is independent of (*A, B*_inc_) and hence of *m*, and *Y* ^*^ = *m* + (*U* − *Û*) + *ε*, giving (i); writing *B*_exc,*j*_ = *ρÛ* + *ρ*(*U* − *Û*) + *η*_*j*_, only the middle term covaries with *Y* ^*^ given *m*, contributing *ρv*, which is (ii) with no Gaussianity of *h*(*A*) needed. *c*^*′*^(*ρ*) = (1 − *kρ*^2^)*/*(1 + *kρ*^2^)^2^ gives the turning point.

The proposition concerns the population covariance with the latent outcome *Y* ^*^; the implemented statistic is the within-stratum adjusted AUC of a cross-fitted probe on the excluded half for the binary *Y* with finite bins, and its positivity at every proxy strength is the observed 100% rejection rate above, not a further theorem.

### Real-descriptor null

For each corpus we keep the 62 real features as group *A* and generate labels from them alone, with log-odds *α* core(*A*) + *u*, where the core is a fixed random linear direction or a nonlinear function of *A*’s first three principal components, *α* ∈ *{*1, 2}, and *u* is standard normal. Nothing the propensity omits drives the label, but the boosted-tree propensity is estimated as in practice, so estimation and functional-form error remain. Over 5 folds × 6 halvings per condition (1200 halvings), the threshold 0.0167 is exceeded in 16.9%: 0–16.7% per corpus for the frequency group and 17.5–33.3% for morphology. This is not the false-positive rate of an exactly correct propensity, and the per-halving threshold does not calibrate a decision based on the median over halvings. A matched calibration of that median under real-descriptor nulls is needed before any real-corpus omission verdict.

### Split stability (descriptive)

Over 50 random halvings per corpus–group case (5 repeated splits × 10 halvings; threshold 0.0167), rejection fractions and median EXC (frequency/morphology) are AFB 0.60/0.72 (+0.0240/+0.0286), malaria 0.28/0.50 (+0.0101/+0.0168), Kaggle 0.48/0.32 (+0.0145/+0.0112), Wellgen 0.36/0.48 (+0.0124/+0.0161), and PCam 0.40/0.36 (+0.0124/+0.0106). Given the null above, these fractions are not evidence of omission; we report them only to document variation across halvings.

### Attribution (exploratory)

Within the propensity strata of group *A*, we score each held-out column by its own within-stratum label signal, 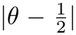 with the column as the score (no fitting), and rank columns by the mean over the five splits. Adding the top three to the 62-feature model moved the zero-shot AFB estimate from 0.767 to 0.766 (frequency) and to 0.759 (morphology); changes on other corpora were at most 0.0051. Baseline and expanded models fail at least one check on AFB, Wellgen and malaria, and ranking and evaluation reused the same split labels, so this is not a validated repair. Whether a ranked column is nuisance or disease signal needs domain input.

## E Exploratory label-fit selection: validation and real-corpus detail

### Rule and validation design

Candidates C1, C1e (linear class), C2, C2r (tree class) on the same 16 features: inside each outer training fold, 4-fold cross-fitting picks each class’s member by log-loss, a paired *z* on the two members’ per-tile log-loss vectors decides the class (|*z*| ≥ 2, otherwise abstain), and the selected member is refit and scored on the test fold, with Brier score as the secondary fit comparison. Validation uses each corpus’s real 16-feature geometry with synthetic labels (as §4) that are linear-in-span, nonlinear-in-span, or outside-span, at two label strengths and audit sizes *n* ∈ *{*full, 1000} with six replicates: 60 conditions, decided by the majority of replicate verdicts when at least half adjudicate, informative when the mean-error gap is ≥ 0.01.

### Validation against ground truth

Table 7 summarises the three validation blocks. Selection never sees the detector score, and on full-size cells with a linear truth the rule chooses the linear class in 238 of 238 decided replicates in the 24-replicate frozen-replication block (two further replicates abstain), so its preference for trees on the real corpora is not a built-in bias. Its downstream benefit is small: the two conditions favouring the linear class in the original block did not retain their original advantage on fresh draws (signed gaps −0.0234 and −0.0133 became +0.0012 and −0.0014), and the new linear-favouring design gains 0.0014 over always choosing trees. Abstention is not fully calibrated: all strongly informative conditions (gap ≥ 0.02) were decided, but the median-ratio calibration criterion failed narrowly (0.562 against ≤ 0.5).

### Selection on real corpora

Table 8 reports the class-selected estimates. The rule also picks trees on Wellgen, *z* 39.7–44.0, and malaria, *z* 4.9–5.9, in every split.

**Table 8:** Label-fit selection on real corpora. Adjusted-AUC pairs are zero-shot/self-trained. Fold counts apply to each detector; *z* ranges span both. The preferred class has lower cross-fitted label log-loss, which does not establish lower adjusted-AUC error.

| corpus | verdict | folds | paired *z* | preferred | disfavored |
|---|---|---|---|---|---|
| AFB | tree | 5/5 | 9.4–16.5 | 0.842/0.865 | 0.890/0.908 |
| Kaggle | tree | 5/5 | 18.8–30.5 | 0.709/0.762 | 0.749/0.807 |
| PCam | tree | 5/5 | 8.5–12.0 | 0.678/0.697 | 0.720/0.750 |

**Table 9:** Adjusted-AUC differences between linear and tree nuisance models, with descriptive 95% cluster-bootstrap ranges. “Selected” uses the selected linear minus selected tree member for zero-shot scores; other columns use fixed C1 minus C2. Self-trained scores come from the transductive student; probes are logistic models on BiomedCLIP or DINOv2 embeddings trained on real labels (raw AUC 0.91–0.99).

| corpus | selected | self-trained | BiomedCLIP | DINOv2 |
|---|---|---|---|---|
| AFB | 0.048 [0.036, 0.060] | 0.051 [0.040, 0.062] | 0.029 [0.023, 0.036] | 0.039 [0.031, 0.047] |
| Kaggle | 0.040 [0.025, 0.051] | 0.050 [0.036, 0.063] | 0.033 [0.026, 0.039] | 0.022 [0.017, 0.028] |
| PCam | 0.042 [0.032, 0.080] | 0.055 [0.047, 0.096] | 0.053 [0.046, 0.085] | 0.029 [0.015, 0.045] |
| Wellgen | 0.048 [0.003, 0.095] | 0.041 [−0.013, 0.092] | 0.117 [0.096, 0.138] | 0.064 [0.049, 0.080] |
| malaria | −0.015 [−0.025, 0.002] | −0.005 [−0.010, 0.007] | 0.003 [−0.001, 0.012] | 0.004 [0.000, 0.012] |

Among the three tabulated corpora, the chosen member is C2 in 22 of 30 fold-selections and C2r in 8; the Brier verdict agrees in 30 of 30. Preferred and disfavored levels are means over folds of each class’s selected member and therefore differ in the third decimal from Table 4’s fixed C1 and C2 (AFB zero-shot 0.890 versus 0.894). On these three corpora the decision does not change for any abstention threshold in [0, 8.5) (on all five, in [0, 4.87)), and on full-size validation cells with a linear truth the original six-replicate block picks the linear class 60 of 60 times. With each corpus’s available unit (Kaggle: 344 filename-derived image groups, largest 70 tiles; DINOv2 near-duplicate clusters at *τ* = 0.95 where no slide identifier ships, AFB: 5502 clusters, largest 182; PCam: 2939 clusters, largest 4902, a dominant component holding 61% of tiles), the unit-level bootstrap 95% interval of the class log-loss gap excludes zero on the tree side in 30 of 30 fold-analyses (five original and five grouped outer folds per corpus), cluster-robust *z* 6.4–19.3; this resamples the log-loss gap, not the paired conditional-AUC difference, and slide structure unmeasured on AFB and PCam remains outside any computable unit.

## F Uncertainty, other detectors, and diagnostics

### Cluster bootstrap

All intervals for C1 minus C2 are paired: both models are evaluated on the same 1000 bootstrap draws of the resampling units of Table 6, with the fitted models held fixed. They therefore measure test-sample variability, not refitting uncertainty or slide and patient dependence the corpora do not record; on PCam one cluster holds 61% of tiles, which also limits the bootstrap approximation. The cluster-bootstrap intervals are empirical bootstrap quantiles. The resampling groups are not established independent sampling units, so the intervals’ population coverage is not established and their exclusion of zero is descriptive rather than a calibrated significance claim. Fold-level paired differences (mean, fold SD) are AFB +0.0553 (0.0063) zero-shot and +0.0505 (0.0062) self-trained, Kaggle +0.0480 (0.0122) and +0.0504 (0.0080), PCam +0.0443 (0.0078) and +0.0550 (0.0100), positive in 5/5 folds in every case.

### Grouped refits and other detectors

With outer folds that keep resampling units apart and refit both models, zero-shot C1 minus C2 is +0.054 on AFB (fold SD 0.012, 5/5 positive), +0.034 on Kaggle (0.011, 5/5), +0.026 on Wellgen (0.039, 4/5) and −0.0125 on malaria (0.036, 2/5); grouped PCam folds contain one label, so their AUCs are undefined. Zero-shot PLIP and QuiltNet scores, whose raw AUCs are 0.30–0.79, give gaps of mixed sign, from −0.050 to +0.105, some with intervals excluding zero.

### Per-fold checks

Table 4 applies the checks to fold-averaged one-label shares and controls. Per fold (zero-shot), C1 passes in 5/5 splits on AFB, Kaggle and Wellgen, 4/5 on PCam and 3/5 on malaria; C2 passes in 5/5 on Kaggle, 4/5 on PCam, 3/5 on AFB (its control leaves the window in two splits) and 1/5 on Wellgen and malaria. Colour-shift checks are counted per fold; direction agreement is counted after averaging over five splits.

### Bin-count sensitivity

Table 10 recomputes the zero-shot comparison with *q* ∈ *{*5, 10, 20, 40} equal-count bins on the same splits; *q* = 20 reproduces Table 4. The gap is positive in 5/5 splits at every *q* on AFB, Kaggle, PCam and Wellgen and negative in 5/5 on malaria; the checks’ per-split pass counts change with *q*.

### Colour-shift design

Each tile receives the shift with probability *p*_1_ if positive and *p*_0_ if negative: (0.5, 0.5) independent, (0.8, 0.2) with the label, (0.2, 0.8) against it; on the four class-balanced corpora every condition shifts half the tiles in expectation (on AFB, 40% and 60% under the label-dependent assignments). The supervised probe is trained on the training fold of the shifted corpus. The detector’s loading on the shift is the paired shifted-minus-original mean score change, scaled by the original-score SD. The zero-shot loading uses all tiles at each shift strength (−0.165 to +0.335); each supervised loading uses only that probe’s test fold, separately for each condition. The reference for C1 or C2 is the pair-weighted AUC within its 20 original-image propensity bins crossed with the shift indicator. Five splits are averaged within each of 80 label-dependent cells (5 corpora, 2 values of *δ*, 2 assignments, 2 detectors, 2 models). The rule chose trees in all 40 corpus–*δ*–assignment–detector conditions.

### Four same-feature learners

(C1e is an elastic-net logistic model and C2r a random forest; mean conditional AUC over the same five folds, zero-shot/self-trained; on AFB, Kaggle and PCam all four pass both checks on fold-averaged values for both scores, though individual splits can fail, as Table 4 shows for C1 and C2): C1e reads 0.890/0.908 on AFB, 0.749/0.807 on Kaggle, and 0.720/0.750 on PCam, and C2r 0.854/0.871, 0.711/0.762, and 0.682/0.698, with C1 and C2 zero-shot as in Table 4 and self-trained 0.914/0.863 on AFB, 0.811/0.761 on Kaggle, and 0.752/0.697 on PCam. Pairwise mean absolute disagreement: within class, C1–C1e 0.003 and C2–C2r 0.006; across classes, 0.041–0.051 for every pair.

### Sensitivity to the adjustment set

The same tree learner on 62 features (C3) reads 0.767/0.790 on AFB, 0.644/0.690 on Kaggle, and 0.626/0.640 on PCam against C2’s 16-feature 0.838/0.863, 0.705/0.761, and 0.677/0.697: the estimate changes by 0.051–0.073 and C3 fails the checks on AFB, a change in what is estimated, accompanied by a failed check. Under the independence assumption of Appendix A the movement may reflect shortcut removal as well as estimation, binning and changes in pair weights; if texture or edge statistics carry disease signal, part of it may be over-adjustment, and the two cannot be separated without labels that distinguish them.

## G Implementation details

### Detectors

The zero-shot detector is BiomedCLIP (Zhang et al., 2023) (checkpoint `microsoft/ BiomedCLIP-PubMedBERT_256-vit_base_patch16_224`, loaded with open_clip and its own image pre-processing). Image embeddings are L2-normalised, and the score of an image with embedding *E* is 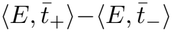, where 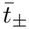 is the unit-normalised mean of the text embeddings of four positive or four negative prompts for the corpus’s stain (Table 11). PLIP (`vinid/plip`) and QuiltNet (`wisdomik/QuiltNet-B-32`) use the same prompts and score. The self-trained student takes the top and bottom 20% of tiles by the zero-shot score as pseudo-labels and fits a standardised logistic regression on the BiomedCLIP embeddings, twice, scoring every tile with a cross-fitted model. Supervised probes are standardised logistic regressions (*C* = 0.1) on BiomedCLIP or DINOv2 embeddings, trained on each split’s training tiles with the real labels. DINOv2 vectors are mean-pooled ViT-S/14 patch tokens, L2-normalised.

**Table 10:** C1 minus C2 (zero-shot, mean over five splits) by number of strata *q*. Parentheses give pass counts for C1/C2, each out of five splits; a pass requires both checks.

| corpus | *q* = 5 | *q* = 10 | *q* = 20 | *q* = 40 |
|---|---|---|---|---|
| AFB | +0.050 (1/0) | +0.053 (4/2) | +0.055 (5/3) | +0.058 (5/2) |
| Kaggle | +0.043 (5/0) | +0.045 (4/0) | +0.048 (5/5) | +0.050 (5/5) |
| PCam | +0.040 (0/0) | +0.046 (2/0) | +0.044 (4/4) | +0.044 (5/4) |
| Wellgen | +0.045 (1/0) | +0.045 (3/0) | +0.046 (5/1) | +0.048 (5/0) |
| malaria | −0.010 (0/0) | −0.014 (0/0) | −0.015 (3/1) | −0.014 (5/1) |

**Table 11:** Zero-shot prompts. Kaggle, AFB and Wellgen use the Ziehl–Neelsen prompts, PCam the H&E prompts, malaria the Giemsa prompts.

| stain | positive | negative |
|---|---|---|
| ZN | a photomicrograph of a Ziehl-Neelsen stained sputum smear containing red rod-shaped acid-fast bacilli<br>microscopy of sputum with red acid-fast bacilli against a blue background<br>Ziehl-Neelsen stain showing Mycobacterium tuberculosis bacilli<br>a sputum smear positive for acid-fast bacilli | a photomicrograph of a Ziehl-Neelsen stained sputum smear with no bacteria<br>microscopy of sputum showing only blue stained background material<br>Ziehl-Neelsen stain with no acid-fast organisms present<br>a sputum smear negative for acid-fast bacilli |
| H&E | a histopathology image of lymph node tissue containing metastatic breast carcinoma<br>H&E stained lymph node with tumor cells<br>microscopy of lymph node showing metastatic tumor<br>a lymph node section positive for metastasis | a histopathology image of normal lymph node tissue<br>H&E stained lymph node without tumor cells<br>microscopy of benign lymph node tissue<br>a lymph node section negative for metastasis |
| Giemsa | a photomicrograph of a Giemsa stained red blood cell infected with a malaria parasite<br>a blood smear showing a red blood cell containing Plasmodium<br>microscopy of a parasitized erythrocyte<br>a thin blood film positive for malaria parasites | a photomicrograph of a normal uninfected red blood cell<br>a blood smear showing healthy erythrocytes<br>microscopy of an uninfected red blood cell<br>a thin blood film negative for malaria parasites |

### Nuisance descriptors

Pixel values are scaled to [0, 1]. The 16 colour and focus features are the mean and standard deviation of seven channels (red, green, blue, HSV saturation and value, and fuchsin and methylene-blue optical densities from a fixed colour-deconvolution matrix) plus the variance of the Laplacian and the grey-level standard deviation. The 62-feature descriptor adds the 1st, 10th, 50th, 90th and 99th percentiles of the red, green, blue and two optical-density channels (25); six grey-level co-occurrence statistics (64 levels, distance 1, four angles averaged: contrast, dissimilarity, homogeneity, energy, correlation, angular second moment); a 10-bin histogram of uniform local binary patterns (*P* = 8, *R* = 1); the mean, standard deviation and 90th percentile of gradient magnitude; the mean absolute Laplacian; and the fraction of pixels with Sobel magnitude above 0.1. Frequency and morphology groups are as described in Appendix C.

### Nuisance models

All learners are from scikit-learn. C1 is a standardised L2 logistic regression (*C* = 1, L-BFGS, up to 5000 iterations). C2, C3 and C4 are histogram gradient-boosted trees (200 iterations, learning rate 0.1, at most 31 leaves, other settings default, seed equal to the split index). C1e is a standardised elastic-net logistic regression (SAGA, *l*_1_ ratio 0.5, *C* = 1); C2r is a random forest (400 trees, at least 20 tiles per leaf).

### Splits, strata and ties

Each of the five splits takes the first fold of a shuffled stratified 4-fold partition with seed *r* ∈ *{*0, …, 4} (75% training, 25% test). Nuisance models are fitted on training tiles and evaluated on test tiles. Bin edges are the empirical *k/q* quantiles of the test-fold propensities, and each tile goes to the bin given by the number of interior edges at or below its propensity. Tied propensities therefore always share a bin, so tree propensities, which take few distinct values, can produce unequal bins. Within each bin the detector’s AUC is computed with scikit-learn’s ROC AUC, which gives tied scores half credit. Bins containing one class contribute nothing and are counted for the overlap screen. Bins are weighted by their number of positive–negative pairs. The self-stratified control applies the same function with the propensity as the score, so its within-bin ties also receive half credit. Both checks are computed per split and, unless stated otherwise, averaged over splits.

### Resampling

Each bootstrap draw resamples the units of Table 6 with replacement once, and the same multiset is used for all splits, both models and all detectors, so differences are paired; tiles enter with their unit’s multiplicity, fitted models are not refitted, and the bracket is the 2.5th–97.5th percentile of 1000 draws. Near-duplicate clusters are connected components of the graph that joins tiles whose DINOv2 vectors have cosine similarity at least 0.95. Grouped refits use stratified group 4-fold partitions (seed 100 + *r*) that keep units apart.

#### Ties and bin masses

On the five corpora’s real-data comparisons (zero-shot detector, *q* = 20), the fitted C1 and C2 propensities have at most 26 tied values among the 2000 test tiles of any split (on PCam; none on AFB), bin sizes differ by at most 1.3%, and no tile lies in a bin more than 25% away from *n/q*. Recomputing with bins weighted by *π*_*k*_(1 − *π*_*k*_) instead of pair counts, or after breaking ties with a random jitter of order 10^−9^ (10 draws), changes no split’s C1 or C2 by more than 0.0002; tie-breaking alone changes nothing.

### Software and data

Python 3.9 with scikit-learn 1.6.1, NumPy 2.0.2, SciPy 1.13.1, PyTorch 2.8.0 and open_clip 3.3.0. What can be reproduced from which inputs is stated in the data and code availability section.

## H Deterministic reruns and historical counts

The archived semi-synthetic generators seeded with Python’s process-dependent hash(corpus), so their draws cannot be reconstructed (Appendix C). We reran the unchanged designs under a deterministic SHA-256 corpus seed. These are new draws, not a reconstruction, and all three-regime grid counts in the paper come from them.

### Historical counts

The archived three-regime grid gave direction agreement in 583 of 600 non-oracle cells (deterministic: 590), reversal of the 16-feature logistic model at *λ* = 3 against the label in 13 of 50 replicates (14) and at *λ* = 2 in 6 of 50 (5), an AFB mean of 0.475 against a truth of 0.554 (0.481 against 0.566), 600 of 600 cells passing the checks (594), and a mean control of 0.513 (0.5105). Its 17 sign disagreements were single replicates in no-loading conditions and in tree-model conditions on AFB and PCam. The qualitative findings agree; the counts depend on the draws.

### Other confirmations

Measurable/residual-driver sweep (750 rows): oracle mean absolute error 0.0040; all 21 linear-16 cells with error above 0.05 pass the screen; at its own calibration threshold 0.01766 the residual driver’s rejection rate is at most 0.10 at every strength while error among non-rejecting cells reaches 0.0696 (95th percentile 0.0557); the measurable-driver criterion (in each corpus at *δ* ≥ 0.5, rejection fraction at least 0.80 or mean error at most 0.03) fails on malaria (rejection 0.00, error 0.0332) while the other four corpora meet the error criterion.

Construction-sensitivity confirmation (450 rows): the oracle is recovered with mean absolute error 0.0037– 0.0043 across the three label directions and fitted-tree error is larger in all three (0.0146–0.0205); at driver strength *δ* = 1 with the within-run threshold 0.0173, only two of the four residual directions meet both rejection fraction ≤ 0.20 and error ≥ 0.03 (rejection fractions 0.08, 0.00, 0.32, 0.32; errors 0.0483, 0.0360, 0.0529, 0.0533), so the criterion requiring three directions fails. Undetected bias remains, but the historical three-of-four robustness result does not replicate; no threshold was retuned.

Regenerated Family I, three-family screen, and real-corpus (Table 4) tables match their historical counter-parts exactly from the cached inputs, and 33 AFB replicate rows replay exactly under a different Python hash seed. The additional experiments (cluster bootstraps, other detectors, colour shift, real-descriptor null, attribution, linear-favouring design, law grid and bins) use fixed seeds.

## I Experiment inventory

Table 12 lists every experiment, the question it answers, and where it is reported. The names in typewriter type are the experiments of the supplementary package; each runs with nuisauc run *name* from its default configuration.

**Table 12:** Experiments in this paper, with the package experiments that produce them.

| experiment | question | where | main result |
|---|---|---|---|
| Gaussian Families I and III, quadrature vs. simulation<br>theory_law, theory_montecarlo, law_grid | Does the formula predict adjusted AUC? | §3, App. B | 36 cells within 0.008 |
| Gaussian Families I–III, checks<br>family_i, families_ii_iii, control_bins, cutoff_sweep, overlap_medmnist | Do the checks catch material errors? | §4 | 140 of 141 material cells pass |
| Semi-synthetic three-regime grid<br>semisynthetic_regimes, oracle_bins | Does the predicted direction hold with practical models on real descriptors? | §3, App. C | 590/600 replicate directions; reversal only under adversarial loading |
| Real-label panel (C1 vs. C2)<br>realdata_panel, realdata_screen, realdata_uncertainty, realdata_enrichment | Do same-feature models agree? | §5 | AFB/Kaggle/PCam: gaps 0.044–0.055; averaged checks pass |
| Bin-count sensitivity<br>bins_balance, ties | Does the gap depend on *q*? | §5, App. F | gap stable; check verdicts vary |
| Fit diagnostics<br>realdata_panel, bins_balance, realdata_learners | Do broader diagnostics separate the models? | §5 | C1 imbalance above the permutation floor |
| Other detectors<br>realdata_panel | Is the gap detector-specific? | §5, App. F | supervised probes 0.022–0.117 |
| Colour shift on real images<br>colour_shift | Does the predicted direction hold for a controlled shortcut? | §5, App. F | 78 of 80 cells |
| Selection-rule validation<br>selection_validation, selection_cluster, selection_replication, selection_linear | Does label fit pick the more accurate class? | §5, App. E | small gains |
| Held-out tests (split-family test)<br>heldout_statistic, heldout_specificity, heldout_hob, heldout_lod, heldout_null, heldout_attribution | Can a held-out group flag omission? | §5, App. D | not calibrated for real descriptors |
| Deterministic reruns<br>semisynthetic_checks, semisynthetic_sensitivity | Are the counts reproducible? | App. H | undetected bias persists; counts and robustness differ |

